# Where do animals covered by the US Animal Welfare Act live? Analysis of USDA licensed and registered entities

**DOI:** 10.1101/379412

**Authors:** Allyson J. Bennett, Erin G. Schoenbeck

## Abstract

Nonhuman animals are housed in captivity for a range of purposes in the US and other countries. The regulation, oversight, and public transparency of decisions related to the care and use of those animals varies by species and activity. The goals of this paper are 1) to provide a concise and accessible summary of the number and types of facilities either registered or licensed by the USDA and overseen by the federal agency; and 2) to provide concise comparisons of the relative proportion of different types of use of nonhuman animals that fall under the AWA (Animal Welfare Act). Analysis of the publicly-available list of USDA certificate holders produced descriptive data for each state and the US overall. Licensed exhibitors (N=2,640; 33% of total) and breeders (N=2,701; 34% of total) comprise two-thirds of the 8,002 USDA certificate holders. Registered research facilities (N=1,110) account for 14% of the total USDA certificates. The final 19% consists of licensed dealers (N=763; 9%) and registered carriers or interim handlers (N=788; 10%). The number and distribution of types of certificates varies across states. The largest number of exhibitors and dealers are in Florida, while the largest number of breeders are in Missouri. California has the largest number of research registered facilities. Finally, a comparison of the estimated number of animals in the US that are pets, used in agriculture, exhibition, and research suggests that animals in research are between 0.07-2.3% of the total. The comparison also highlights areas in which public information about captive animals in the US is uneven or missing altogether. Together, the current report provides a concise view of basic information and data relevant to public consideration and policy decisions about animals housed in the set of captive settings to which the AWA applies.

## Introduction

Nonhuman animals are housed in captivity for a range of purposes in the US and other countries. For some of those animals, federal legislation governs their treatment and standards of care, including provisions for external oversight by federal agencies. As a result of this mandated federal oversight there are also mechanisms for public transparency about decisions related to the acquisition, production, maintenance, transfer, and use of those animals. Regulatory protection, oversight, and public transparency vary by species, the purpose (or activity) for which the animals are in captive settings, and the type and location of the facility, or setting, in which they are housed (for review see reference 4).

The 1966 Animal Welfare Act (AWA) and subsequent amendments provides one of the main sources of consistency in federal regulation, external oversight, and public transparency related to captive animal care.^2,3^ The United States Department of Agriculture (USDA) is the federal agency charged with information collection and systematic monitoring relevant to the AWA, as well as additional federal oversight. The AWA applies only to a subset of captive animals and excludes the large proportions that are used in agriculture, as companions (pets), and for some other purposes. Further, it applies only to a subset of species (see detail below). Nonetheless, it is the major venue by which information is collected, care standards promulgated, and external oversight applied over a diverse range of facilities and species housed in captive settings.

Provisions of the AWA both directly and indirectly affect the care, oversight, and availability of public information about captive animals. The USDA provides licensure or registration to entities engaged in the exhibition, breeding, sale, transport, handling, research, or testing of a broad range of nonhuman animals. Together, licensees and registrants are termed certificate holders. Figure 1 describes the different types of USDA certificates. However, there are few sources for concise demographic information about the entities overseen by the federal agency, and broad comparison of the number and type of regulated entities is not readily available in a concise and accessible format. Moreover, for species to which the AWA, applies as well as those not covered by federal legislation, there is little consistent information about the number and type of animals housed in various captive settings in the US.

**Figure 1.**
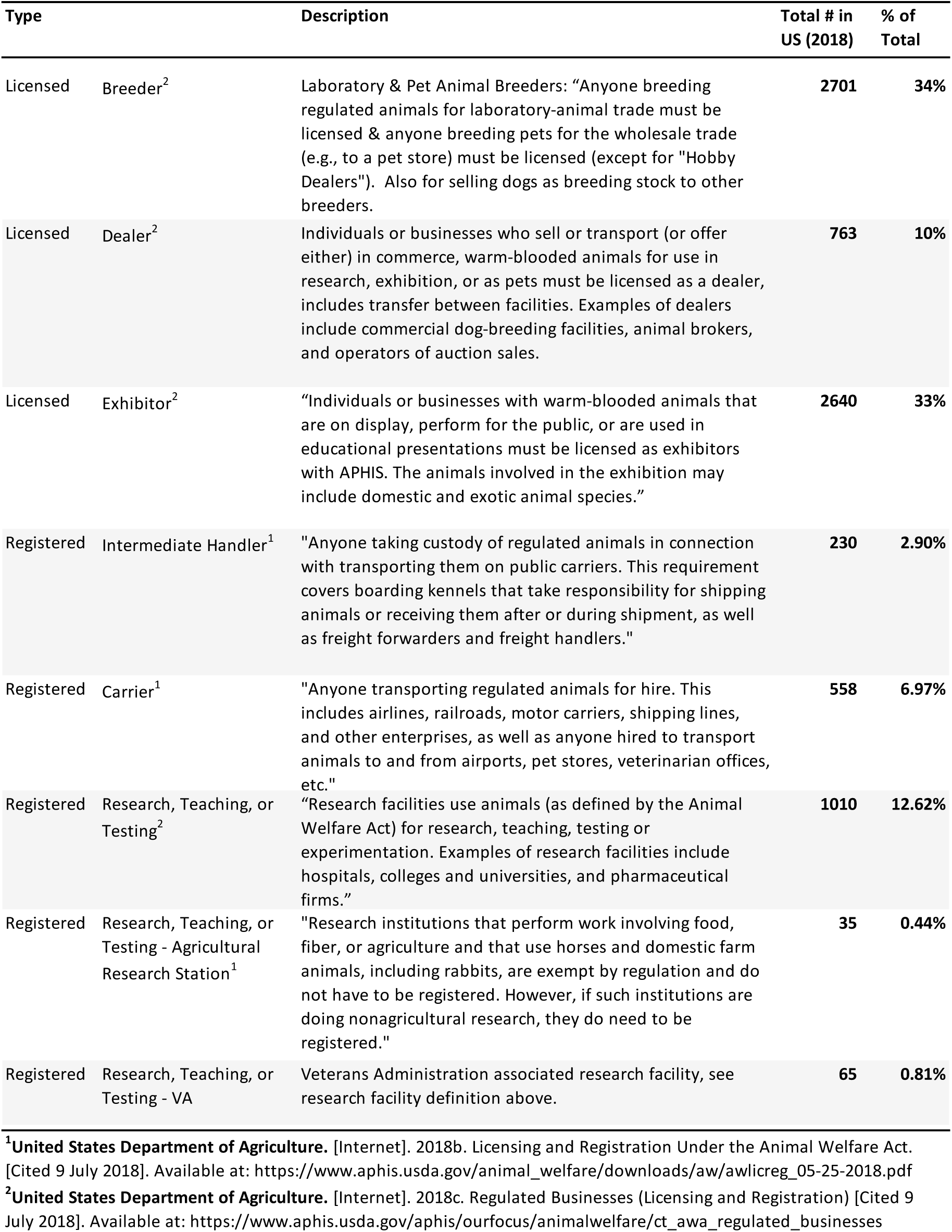
Overview and description of USDA license and registration categories.

Accessible basic demographic information about the composition and number of entities licensed or registered to be overseen by the USDA for compliance with the AWA is important for many reasons. Among them, it provides critical context for decision-making, policy changes, and public understanding of the use of captive animals. Information about the relative proportion of animals in different types of facilities, or falling under different types of regulatory oversight, is important to inform policy related to animal care and welfare. Such information can also be applied to help understand the relative costs and benefits of proposed legislative or policy changes regarding oversight and public transparency of facilities. More broadly, the quality of critical assessment and weighing of the impact of proposals for changes in any of the standards for animal care depends on foundational data about the composition and animal numbers within the full range of facilities and entities that fall under AWA and USDA oversight.

Full demographic data about USDA certificate holders is also the requisite foundation for accurate description and context from which to consider the number of animals in research, testing, and education. In the US, but particularly in Europe, the number of animals used in research and testing receives public attention and frequent news coverage. The rationale for focusing on animal use numbers is often explained in terms of providing the public with important data that can inform their perspectives, understanding, and decisions about the use of animals in research and testing. Further, these data are often compared across time and across countries. Such comparisons are often cast in terms of monitoring adherence to a simple heuristic often referred to as capturing the ethical considerations in animal research, the “3 Rs”.^19^ In this heuristic, one goal is to reduce the number of animals used to answer scientific questions or to test medical products. The other two “Rs” call for replacement of live animals with alternative methods (e.g., simulations, cell culture, etc.) and refinement of methods in order to increase animal welfare.

In contrast to replacement and refinement, the number of animals is easily quantified. It is perhaps for this reason that the number of animals in research is a focus of regulatory requirement and public reporting. The reporting and accompanying attention often ignore the broader context: the proportion of animals in research relative to the total number of animals used by humans in a variety of other activities. The range of activities include food consumption, manufacture of clothing and other consumer goods, companionship, entertainment, education, commercial promotions, and labor. The absence of the broader context poses challenges to reliable comparisons across time and across countries. For instance, it is likely that the number of animals in captivity varies with the size of the human population. The number of animals, or the relative number of facilities for a particular activity, is also likely to vary in an orderly fashion with such factors as size of the economy; primary industries including agriculture and tourism; relative strength of the educational and biotechnology sector; and so on. To our knowledge, there is little explicit analysis of the factors that influence changes in the number of animals or the proportion of animals involved in various human activities. As a result, there is little information from which to guide estimates of the broad impact of policy or practice changes either in the immediate term or in the future.

### Types of AWA-regulated entities

In the US, as in other countries, a large proportion of animals bred and maintained in captive settings fall under the broad umbrella of agricultural use. A range of species including cows, goats, pigs, chickens, and other animals are used for food production, material for clothing, furniture, and other consumer goods. Another broad category and large proportion are those used for human companionship and for human entertainment. Companion and entertainment animals are predominantly domesticated species that include dogs, cats, guinea pigs, hamsters, etc. but also a wide range of other species that extends into non-domesticated, “exotic” animals. Captive animals are also housed within facilities that provide for exhibition, including those whose goal is public education, or support for conservation of the species, or both. Finally, a relatively smaller proportion of animals in captivity are used for scientific research, and for testing drugs, devices, and products used by humans, but also for the treatment and care of other animals.

Whether an animal falls under the AWA depends both on its species and its use (or activity). According to the USDA:

The following animals are not covered: farm animals used for food or fiber (fur, hide, etc.); cold-blooded species (amphibians and reptiles); horses not used for research purposes; fish; invertebrates (crustaceans, insects, etc.); or birds, rats of the genus *Rattus*, and mice of the genus *Mus* that are bred for use in research. Birds (other than those bred for research) are covered under the AWA, but the regulatory standards have not yet been established.^9^

Note, however, that the very same species may not be included in AWA coverage because inclusion also depends upon the purpose for which the animal is housed in captivity. In short, the law applies to “certain animals exhibited to the public, sold for use as pets, used in research, or transported commercially.”

As evident from the list of animals and activities excluded from AWA and USDA coverage, one of the major divisions is between farm animals (i.e., agricultural use, food, fiber) and those used for other purposes. Thus, a cow that is involved in a *research* study falls under AWA and USDA. In contrast, a cow bred and maintained for milk or meat consumption, but not involved in research, is *not* covered by the AWA, but likely falls under other types of federal, state, or local regulatory oversight. Similarly, privately-owned animals that are not exhibited or involved in large-scale commercial ventures (i.e., public contact or commerce) are not covered by AWA or USDA oversight. Thus, a dog in research, in a commercial breeding and sales facility, or in some zoos or entertainment facilities would be covered, while a pet dog or a dog in a small direct-sale breeding operation (less than 7 dogs) would not be covered. Again, however, as in the example of the cow above, the pet dog or a dog in a non-AWA activity may be covered by other laws and oversight.

For species and activities covered by AWA, the USDA may issue a license or a registration. Licenses are grouped into the following major categories, as shown and defined in Figure 1: exhibitors, dealers, and breeders. The category “exhibitor” is an umbrella for a broad range of facility types that include a large variety of entertainment and education providers-- “circuses, zoos, educational displays, petting farms/zoos, animal acts, wildlife parks, marine mammal parks, and some sanctuaries.” The definition of exhibitor is: “Individuals or businesses with warm-blooded animals that are on display, perform for the public, or are used in educational presentations.” Commercial dealers are those who “sell or offer to sell or transport or offer for transportation in commerce, warm-blooded animals for use in research, exhibition or as pets.” Licensed dealers are those who sell or broker the sale of animals, but do not breed them, while licensed breeders are those who own a colony and breed them for sale. Finally, USDA registration is required for individuals commercially transporting animals. A single entity or facility can hold more than one type of certificate. It should be noted that for the broad categories described here there are many subcategories and cases in which individuals and facilities are exempted from requirement for USDA licensure or registration. For instance, retail pet stores, public pounds, hobby dealers, and “private zoos” in which animals are not displayed to the public are all exempted from license, registration, and oversight. Discussion of each subcategory and their definitions is beyond the scope of the current report (see references 10 and 11 for further detail).

In addition to USDA licenses for exhibition, breeding, and dealing, there are also USDA research-registered facilities that are subject to the provisions of the AWA and are overseen by the federal agency. Entities engaged in research, teaching, testing, or experimentation with regulated species (see above) are required to obtain, and maintain, such registration. Research facilities include a broad range of public and private universities, hospitals, pharmaceutical companies, animal medicine and product companies, and contract research organizations. While the AWA specifies common standards for the care and housing of nonhuman animals in captive settings, there are differences between the practices, policies, features of oversight, and public transparency applied to licensed and research-registered facilities and individuals.

Figure 2 illustrates the major components of federal oversight and associated mechanisms for public transparency that result from the AWA, with specific focus on comparison of research facilities with other certificate holders. The major comparisons are between federal and privately-funded research; USDA licensed exhibitors and dealers; and, finally, activities that are not required to hold a USDA certificate (i.e., private exhibitors, sanctuaries, small breeders, private owners). As illustrated, for all activities and species covered by the AWA, there is a requirement to register or be licensed with the USDA. Certificate holders are subject to some forms of public transparency because the USDA website posts information about the licensees or registrants. Indeed, the analysis and data in the current paper were derived from the USDA website and freely-available information.

**Figure 2.**
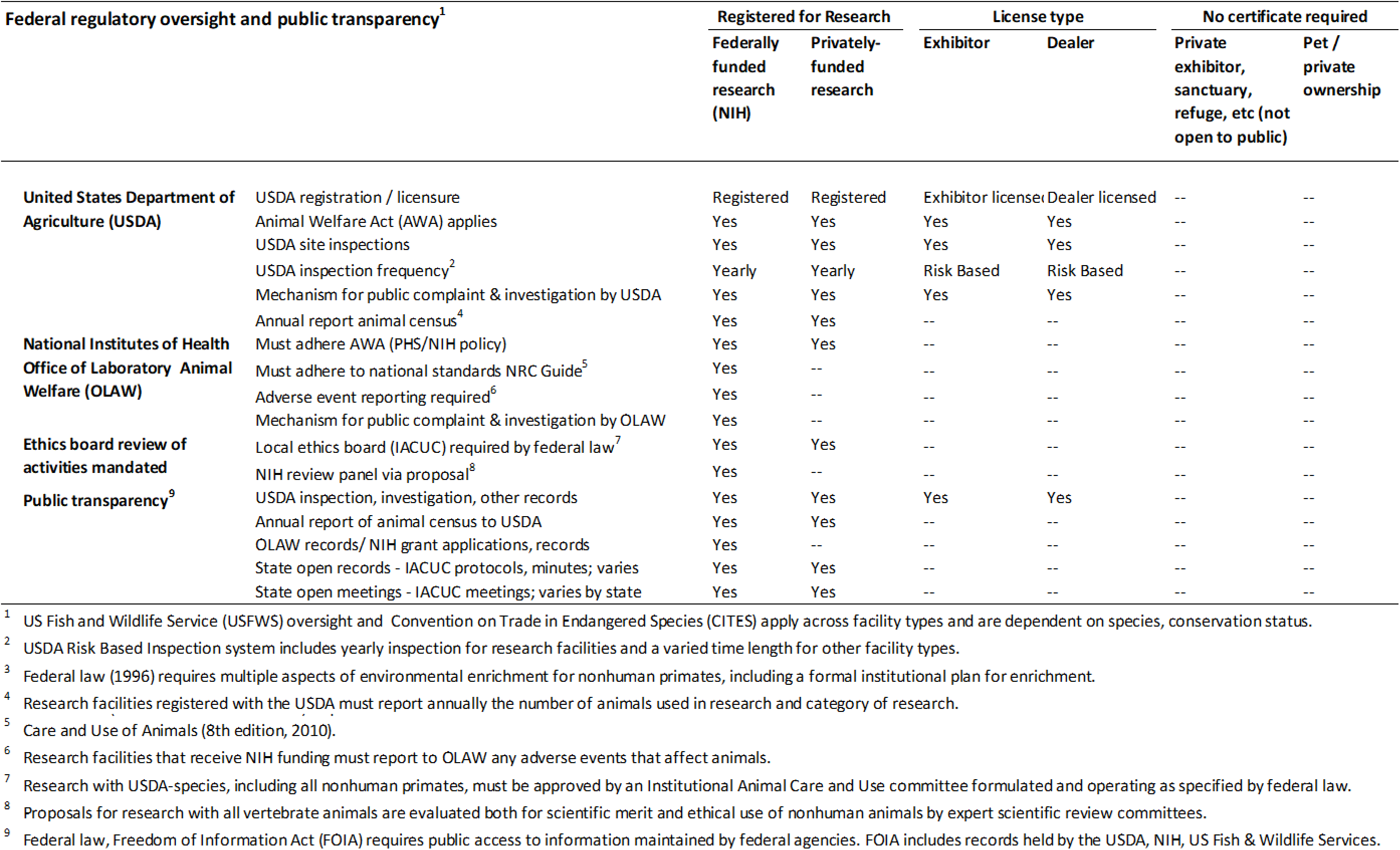
Comparison of regulatory standards, external oversight, and public transparency of different USDA license and registration categories for AWA-covered species.

All certificate holders must also comply with the relevant parts of the AWA and associated standards that apply to their type of activity and the species they hold. Further, there is a mechanism for public complaint to the USDA, which includes potential for the federal agency to investigate the complaint and issue a citation to the certificate holder, as well as an order to correct any problem. The agency may also issue a fine or take action against the certificate holder. The inspection reports are a matter of public record and provide another venue for public transparency.

Along with some common features, there are also significant differences in how the AWA and other federal laws apply to research registrants versus licensed exhibitors and dealers. Primary among them is the fact that while research registrants are inspected by USDA Veterinary Medical Officers annually, there is no such requirement for routine inspections of licensed exhibitors or dealers. Research facilities are required to submit an annual census of the number of animals in their facility to the USDA and that information is made publicly available via the USDA website. There is no such requirement or public posting for other certificate holders. Because the license fees are based in the certificate holder’s revenue gained via the animal activity, exhibitors, breeders, and dealers are required to submit information that is relevant to the number of animals. While this information falls under the federal Freedom of Information Act and can be requested by the public, it is not routinely available in the manner of annual animal number reports for research facilities.^6^ It is also true that animal census for a certificate holder can be included in an inspection report that is posted on the USDA website. Given that there is no requirement for routine inspections, however, using the census information from publicly-available inspection reports is unlikely to provide reliable estimates of animal numbers across the various non-research, certificate holders.

In addition to variation in oversight and elements of public transparency about animal care, Figure 2 also highlights core differences in processes for ethical consideration and decisions about animal treatment and care between certificate holders. For instance, research facilities have a formal process for decisions about animal activities that includes a requirement to submit a detailed proposal for consideration by an Institutional Animal Care and Use Committee (IACUC) comprised of at least one veterinarian, one scientist, and one member unaffiliated with the institution. By contrast, there is no mandated, formal, or public transparency process for the use of animals in exhibition, breeding, or dealing (for detail and additional discussion see reference 4). Federally-funded research with nonhuman animals is subject to additional standards, oversight, and mechanisms for public transparency, and this coverage extends beyond AWA covered species to include purpose-bred rats, mice, and birds (for additional detail, see reference 8). Figure 2 highlights the additional provisions for research animals and contrasts that with animals in exhibition, dealing, and breeding.

The overall goals of this paper are: First, to provide a concise and accessible summary of the number and type of certificates either registered or licensed by the USDA and overseen by the federal agency, these data are provided for each US state and territory. Second, to provide concise comparisons of the relative proportion of different types of use of nonhuman animals that fall under the AWA. Together, the overarching goal is to provide a concise view of some of the basic information and data relevant to public and policy consideration of animals housed in the set of captive settings to which the AWA applies.

## Method

Data on USDA certificate holders was obtained by accessing the publicly-available list of all active licensees and registrants that is posted on the USDA website. The list was downloaded June 1, 2018 and provides a complete roster of US certificate holders in each of the US states and territories at that time.^12^ It is important to note that the website list is updated frequently and hence the current analysis is a snapshot reflective of data available on the date of download. The list contains: the certificate/customer type (license or registrant), the renewal date, the legal name of the entity, the DBA (Doing Business As) name, its city, and state. The document was transposed into an excel file for further analysis.

The USDA issues three types of licenses: breeder, dealer, and exhibitor. There are four types of registrants: carrier, intermediate handler, research facility, and Veterans Administration (VA) hospital. Figure 1 contains the definition for each. For analysis, the total number of each type of certificate holder and overall total of certificates was calculated for each state and territory. From these totals, the minimum, maximum, average, and standard deviation for both the overall total and each certificate type were calculated for the US states. Descriptive data for the three territories listed with certificate holders was calculated separately given the comparatively low number of certificate holders. Finally, the proportion of certificate types was calculated for each state and territory.

## Results

A total of 8,002 US and territorial entities with species and activities covered by the AWA were licensed or registered for research with the USDA as of June 1, 2018. The category with the largest number of certificate holders was breeders, with a total of 2,701 entities licensed by the USDA (34% of the total). The second largest category was licensed exhibitors, with a total of 2,640 entities (33% of the total). Together, licensed breeders and exhibitors made up the majority, just over two-thirds, of all AWA-covered USDA licensed and registered entities. As illustrated in Figure 3, the 1,110 facilities registered for research with AWA-covered species comprised 14% of the total. The final 19% consisted of roughly equal numbers of licensed dealers (*N*=763; 9% of the total) and registered carriers or intermediate handlers (*N*=788; 10% of the total).

**Figure 3.**
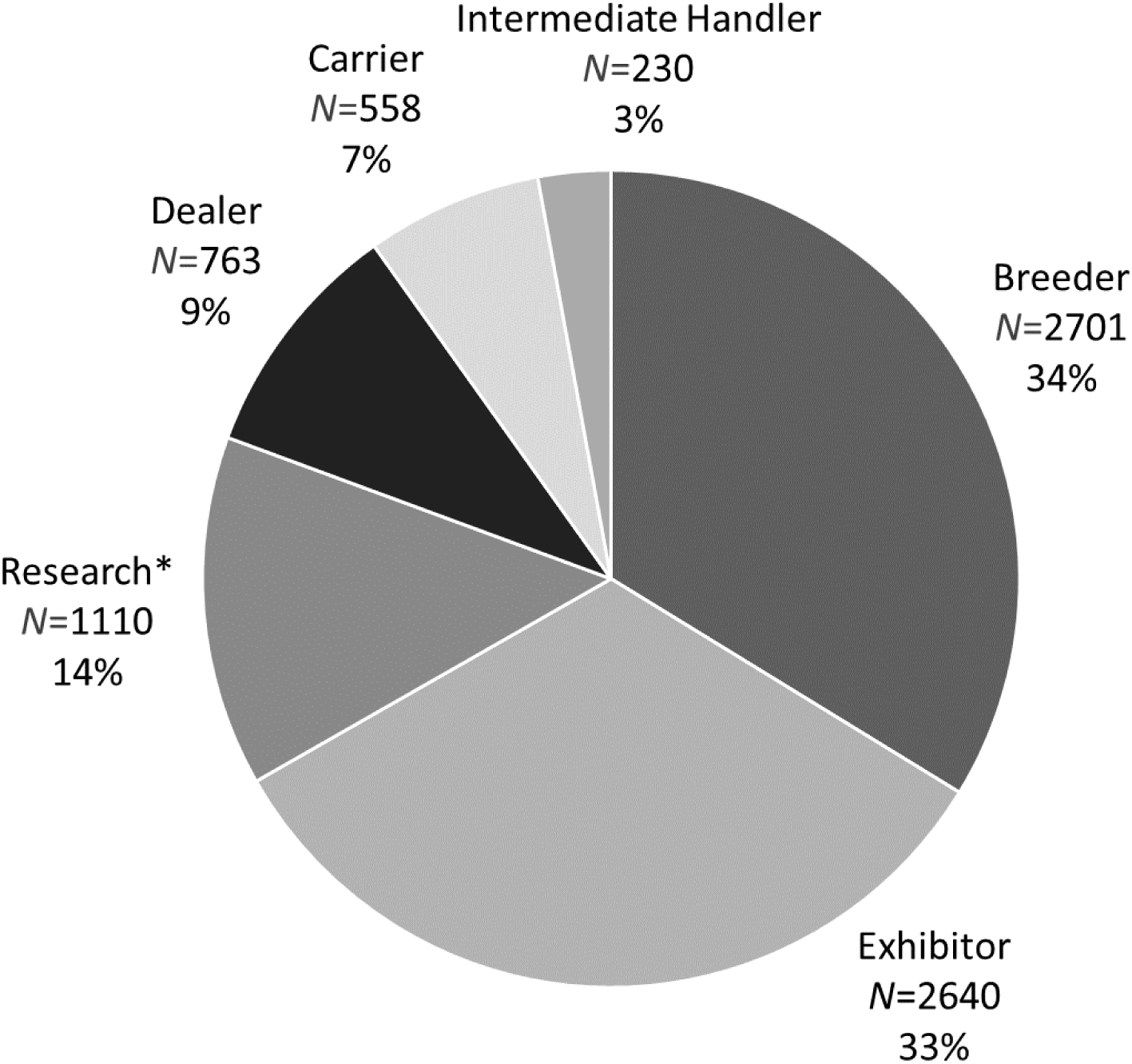
Number and percentage of USDA certificates by category for the United States and its territories. The six major categories of certificates shown include: breeder, exhibitor, research, dealer, carrier, and intermediate handler. Registered research facilities, agricultural research facilities, and Veterans Administration research facilities are combined in this illustration.

### Overall distribution of USDA-licensed or registered entities across states

The number, distribution, and percent of entities within each license and registration type varied widely across the 50 US states, the District of Columbia (DC), and three territories for which certificates appear in the USDA list. In the following section we have focused on the 50 US states and DC, the three territories are discussed in the section titled “Territories.” It is important to note, however, that some facilities that are federally-owned do not hold USDA certificates, including, for example the National Zoo in DC and the National Institutes of Health (NIH) in Maryland. Table 1 contains data at the level of state and shows combined data. Figure 4 illustrates the proportion of certificate type at the level of the state, highlighting the six states that hold the largest number of licensed and registered facilities (i.e., Missouri, Florida, Texas, Ohio, California, and New York). The range in number of entities holding USDA license or registration was a minimum of 10 (Delaware) to a maximum of 984 (Missouri). The average number was 156.51 entities (± SD=176.70). Following Missouri, the next five states with the largest number of entities were: Florida (*N*=474), Texas (*N*=465), Ohio (*N*=455), and California (*N*=435). Following Delaware, those with the fewest were: the District of Columbia (*N*=11), Vermont (*N*=11), Wyoming (*N*=11), Rhode Island (*N*=19), and West Virginia (*N*=19).

**Table 1.** Demographics for USDA licenses and registrations by state for the US and territories

| STATE | Type of License |  |  |  | Type of Registration |  |
| --- | --- | --- | --- | --- | --- | --- |
|  | Total and Percent of Total |  |  |  | Total and Percent of Total |  |
|  | TOTAL<br># | Breeder | Dealer | Exhibitor | Total<br>research &<br>VA | Total R<br>carrier & IH |
| Alabama | 71 | 7 (10%) | 4 (6%) | 40 (56%) | 9 (13%) | 11 (15%) |
| Alaska | 22 | 0 (0%) | 0 (0%) | 15 (68%) | 3 (14%) | 4 (18%) |
| Arkansas | 186 | 120 (65%) | 19 (10%) | 19 (10%) | 16 (9%) | 12 (6%) |
| Arizona | 71 | 8 (11%) | 8 (11%) | 29 (41%) | 12 (17%) | 14 (20%) |
| California | 435 | 8 (2%) | 21 (5%) | 227 (52%) | 128 (29%) | 51 (12%) |
| Colorado | 111 | 12 (11%) | 13 (12%) | 43 (39%) | 24 (22%) | 19 (17%) |
| Connecticut | 88 | 4 (5%) | 4 (5%) | 54 (61%) | 16 (18%) | 10 (11%) |
| DC | 11 | 0 (0%) | 0 (0%) | 0 (0%) | 7 (64%) | 4 (36%) |
| Delaware | 10 | 0 (0%) | 2 (20%) | 3 (30%) | 4 (40%) | 1 (10%) |
| Florida | 474 | 37 (8%) | 81 (17%) | 248 (52%) | 29 (6%) | 79 (17%) |
| Georgia | 150 | 17 (11%) | 13 (9%) | 70 (47%) | 20 (13%) | 30 (20%) |
| Hawaii | 30 | 0 (0%) | 0 (0%) | 11 (37%) | 3 (10%) | 16 (53%) |
| Iowa | 323 | 222 (69%) | 35 (11%) | 34 (11%) | 21 (7%) | 11 (3%) |
| Idaho | 26 | 2 (8%) | 1 (4%) | 14 (54%) | 7 (27%) | 2 (8%) |
| Illinois | 249 | 51 (20%) | 19 (8%) | 117 (47%) | 32 (13%) | 30 (12%) |
| Indiana | 390 | 280 (72%) | 17 (4%) | 60 (15%) | 18 (5%) | 15 (4%) |
| Kansas | 224 | 134 (60%) | 34 (15%) | 32 (14%) | 17 (8%) | 7 (3%) |
| Kentucky | 57 | 5 (9%) | 5 (9%) | 27 (47%) | 9 (16%) | 11 (19%) |
| Louisiana | 80 | 18 (23%) | 5 (6%) | 36 (45%) | 14 (18%) | 7 (9%) |
| Massachusetts | 137 | 5 (4%) | 12 (9%) | 46 (34%) | 67 (49%) | 7 (5%) |
| Maryland | 96 | 8 (8%) | 5 (5%) | 36 (38%) | 38 (40%) | 9 (9%) |
| Maine | 19 | 0 (0%) | 1 (5%) | 8 (42%) | 8 (42%) | 2 (11%) |
| Michigan | 197 | 19 (10%) | 18 (9%) | 108 (55%) | 31 (16%) | 21 (11%) |
| Minnesota | 163 | 30 (18%) | 21 (13%) | 72 (44%) | 28 (17%) | 12 (7%) |
| Missouri | 984 | 752 (76%) | 77 (8%) | 73 (7%) | 27 (3%) | 55 (6%) |
| Mississippi | 49 | 9 (18%) | 4 (8%) | 19 (39%) | 9 (18%) | 8 (16%) |
| Montana | 23 | 5 (22%) | 3 (13%) | 11 (48%) | 3 (13%) | 1 (4%) |
| North Carolina | 147 | 10 (7%) | 19 (13%) | 69 (47%) | 27 (18%) | 22 (15%) |
| North Dakota | 27 | 6 (22%) | 3 (11%) | 13 (48%) | 4 (15%) | 1 (4%) |
| Nebraska | 91 | 51 (56%) | 8 (9%) | 19 (21%) | 11 (12%) | 2 (2%) |

Table 1 con't. Demographics for USDA licenses and registrations by state for the US and territories
| STATE | TOTAL<br># | Type of License<br>Total and Percent of Total |  |  | Type of Registration<br>Total and Percent of Total |  |
| --- | --- | --- | --- | --- | --- | --- |
|  |  | Breeder | Dealer | Exhibitor | Total<br>research &<br>VA | Total R<br>carrier & IH |
| New Hampshire | 25 | 5 (20%) | 1 (4%) | 12 (48%) | 2 (8%) | 5 (20%) |
| New Jersey | 108 | 1 (1%) | 14 (13%) | 55 (51%) | 19 (18%) | 19 (18%) |
| New Mexico | 34 | 2 (6%) | 5 (15%) | 14 (41%) | 10 (29%) | 3 (9%) |
| Nevada | 52 | 3 (6%) | 3 (6%) | 33 (63%) | 5 (10%) | 8 (15%) |
| New York | 340 | 41 (12%) | 20 (6%) | 143 (42%) | 84 (25%) | 52 (15%) |
| Ohio | 455 | 291 (64%) | 26 (6%) | 70 (15%) | 36 (8%) | 32 (7%) |
| Oklahoma | 263 | 174 (66%) | 21 (8%) | 47 (18%) | 13 (5%) | 8 (3%) |
| Oregon | 75 | 12 (16%) | 10 (13%) | 35 (47%) | 10 (13%) | 8 (11%) |
| Pennsylvania | 338 | 114 (34%) | 37 (11%) | 109 (32%) | 54 (16%) | 24 (7%) |
| Rhode Island | 19 | 1 (5%) | 1 (5%) | 7 (37%) | 7 (37%) | 3 (16%) |
| South Carolina | 93 | 8 (9%) | 11 (12%) | 44 (47%) | 14 (15%) | 16 (17%) |
| South Dakota | 76 | 45 (59%) | 11 (14%) | 13 (17%) | 7 (9%) | 0 (0%) |
| Tennessee | 148 | 11 (7%) | 23 (16%) | 67 (45%) | 25 (17%) | 22 (15%) |
| Texas | 465 | 64 (14%) | 70 (15%) | 194 (42%) | 82 (18%) | 55 (12%) |
| Utah | 44 | 5 (11%) | 5 (11%) | 20 (45%) | 8 (18%) | 6 (14%) |
| Virginia | 138 | 14 (10%) | 14 (10%) | 69 (50%) | 26 (19%) | 15 (11%) |
| Vermont | 11 | 1 (9%) | 1 (9%) | 2 (18%) | 5 (45%) | 2 (18%) |
| Washington | 98 | 16 (16%) | 9 (9%) | 30 (31%) | 24 (24%) | 19 (19%) |
| Wisconsin | 229 | 70 (31%) | 25 (11%) | 103 (45%) | 22 (10%) | 9 (4%) |
| West Virginia | 19 | 1 (5%) | 2 (11%) | 5 (26%) | 8 (42%) | 3 (16%) |
| Wyoming | 11 | 2 (18%) | 0 (0%) | 3 (27%) | 3 (27%) | 3 (27%) |
| **Total** | **7982** | **2701** | **761** | **2628** | **1106** | **786** |
| **Average** | **157** | **53 (21%)** | **15 (9%)** | **52 (38%)** | **22 (20%)** | **15 (13%)** |
| **Std. Dev.** | **177** | **120 (22%)** | **18 (5%)** | **54 (16%)** | **24 (13%)** | **17 (9%)** |
| **Minimum** | **10** | **0 (0%)** | **0 (0%)** | **0 (0%)** | **2 (3%)** | **0 (0%)** |
| **Maximum** | **984** | **752 (76%)** | **81 (20%)** | **248 (68%)** | **128 (64%)** | **79 (53%)** |

**Figure 4.**
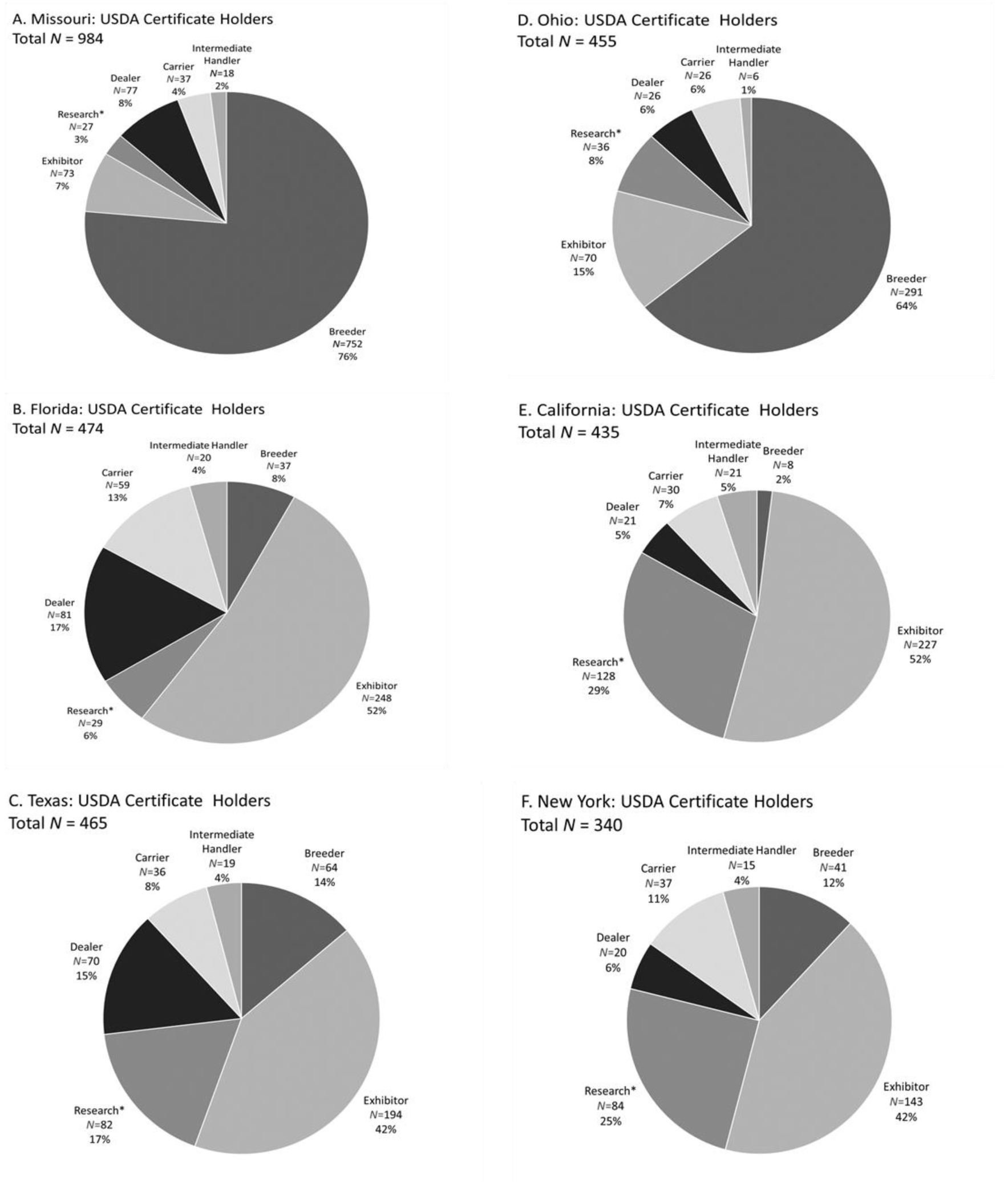
Number and percentage of USDA certificates by category for the six states with the top number of certificates in each major category. A) Missouri is the state with the largest number of certificates (N=984) and largest number of breeders (N=752). B) Florida has the most exhibitors of all the states (N=248). C) Texas has the third highest number of exhibitor licenses (N=194), making up 42% of the total certificates. D) Ohio has the second highest number of breeder certificates. E) California holds the highest number of research registrants. F) New York holds the second most research facilities. Registered research facilities, agricultural research facilities, and Veterans Administration research facilities are combined in this illustration.

The relative proportion of breeders, dealers, exhibitors, carriers, and research facilities varied across states. There were some common patterns, however. For most states, in parallel to the national demographics, the majority of AWA-regulated entities within the state were licensed breeders and exhibitors, with breeders comprising an average of 20.45% and exhibitors an average of 38%, for a combined average of roughly 60%. There were no states in which there were more research-registered facilities than there were entities licensed for breeding, dealing, or exhibition. Within states, research registered entities made up an average of 19.65% (± SD 23.61) of the licensed and registered entities, with a minimum of two and a maximum of 128. In only 13 states did research registration comprise greater than 25% of the total registered or licensed entities.

The proportion of licenses and registrations for Missouri, Florida, Texas, Ohio, California, and New York, highlighted in Figure 4, provide an illustration of variability across states. These six states are those with the highest number of licensed breeders, dealers, exhibitors, and registered research facilities. The total number for each of these is one or more standard deviations greater than the mean across the states.

### Distribution of certificate type across states

#### Breeders

There was a total of 2,701 breeders in the US, making it the largest category of USDA facilities. With respect to license holders for breeding, Missouri, with 752 licensed breeders, was roughly six standard deviations higher than the state average of 52.96 (± SD = 119.90). The next five with the highest number of licensed breeders were also located in central US regions, including Ohio (*N*=291), Indiana (*N*=280), Iowa (*N*=222), Kansas (*N*=134), and Arkansas (*N*=120). By contrast, multiple states had no licensed breeders (Alaska, Delaware, Hawaii, Maine), or only one (New Jersey, Rhode Island, Vermont, West Virginia).

#### Dealers

Florida had both the highest number of licensed dealers and licensed exhibitors. The entire US had a total of 761 licensed animal dealers. Florida’s 81 licensed dealers exceeded the national average of 14.92 by approximately four standard deviations (± SD = 18.08). The next highest numbers were in Missouri (*N*=77), Texas (*N*=70), Pennsylvania (*N*=37), Iowa (*N*=35), and Kansas (*N*=34). In parallel to the state demographics for breeders, a number of states had no USDA-licensed dealers (Alaska, District of Columbia, Hawaii, Wyoming) and several had only one or two (Delaware, Idaho, Maine, New Hampshire, Rhode Island, Vermont, West Virginia).

#### Exhibitors

There was a national total of 2,628 exhibitors in the US. The highest number of exhibitor licenses was found in Florida, where the state’s 248 exhibitor licenses exceeded the national average of 51.53 by almost four standard deviations (± SD=53.76). In contrast to the pattern of a larger number of breeders and dealers in non-coastal, largely Midwestern states, states with the largest number of exhibitors were: Florida (*N*=248), California (*N*=227), Texas (*N*=194), New York (*N*=143), Illinois (*N*=117), Pennsylvania (*N*=109), Michigan (*N*=108), and Wisconsin (*N*=103). These eight states were the only ones with greater than 100 exhibitor-licensed facilities. All of the states had at least two licensed exhibitors. States with the lowest number of exhibitors include: Vermont, with two, and Delaware and Wyoming, with three each.

#### Intermediate Handler

California had the highest number of registered intermediate handlers with 21, out of a US total of 229, which exceeded the national average of 4.49 by roughly three standard deviations (± SD=5.53). The states with the largest number of intermediate handlers were scattered across the US, and included: Florida (*N*=20), Texas (*N*=19), Missouri (*N*=18), New York (*N*=15), Hawaii (*N*=12), and Illinois (*N*=10). These seven states made up the only ones with ten or more intermediate handlers. Several states and DC had no intermediate handlers, including: Delaware, Maine, Mississippi, Montana, North Dakota, Nebraska, Rhode Island, South Dakota, Vermont, and Wyoming.

#### Carriers

There was a total of 557 USDA-registered carriers in the US. Florida contributed the most to this category with 59 carriers. Florida was approximately four standard deviations higher than the US state mean (10.92, ± SD=11.74). After Florida, the states with the largest numbers of carriers were: Missouri (*N*=37), New York (*N*=37), Texas (*N*=36), California (*N*=30), and Ohio (*N*=26). One state had no transporters (South Dakota) and several others had only one (Delaware, Idaho, Montana, North Dakota).

#### Research

Across the states there were 1,041 registered research facilities (non-VA), with an average of 20.41 (± SD=22.36). All of the states had research facilities registered with the USDA. California led with 120, 4.5 standard deviations greater than the national average. California was followed by New York (*N*=79), Texas (*N*=78), Massachusetts (*N*=64), and Pennsylvania (*N*=52). States with five or less registered research facilities included: Alaska, Delaware, Hawaii, Montana, North Dakota, New Hampshire, Nevada, Vermont, and Wyoming.

As discussed above, there are multiple types of research-registered facilities. The majority of states had one or more registered-research facilities associated with the VA. In 24 states there was one such facility, in 10 states there were two-three. Four states had a greater number, with eight in California, five in New York, and four in Texas. States without VA research facilities are shown in Table 1 and largely correspond to states with a relatively small number of USDA research-registered facilities overall.

#### Territories

The US has 14 territories, only three of which had entities listed as licensees or registrants. Together, Guam, Puerto Rico, and the Virgin Islands had 20 USDA certificate holders, with a minimum of one (Guam) and a maximum of 16 (Puerto Rico). There were no breeding facilities and VA research facilities listed as certificate holders in the US territories. Puerto Rico was the only territory with licensed dealers (*N*=2), registered carriers (*N*=1), registered intermediate handlers (*N*=1), and registered research facilities (*N*=4). All the US territories had licensed exhibitors, with the largest number in Puerto Rico (*N*=8). For Guam (*N*=3) and the Virgin Islands (*N*=1), exhibitors comprised all of the territories’ USDA certified facilities.

## Discussion

Comparison of the different types of USDA license holders and registrants provides a broad context for a better understanding of the proportion of activities involving animals covered by the AWA, as well as their distribution across the states and territories. Nationally, exhibitors and breeders make up the large majority, accounting for roughly two-thirds of all USDA certificates. Registered research facilities account for roughly 14% of all certificates. Dealers comprise 9% of certificates. The remaining 10% are carriers and intermediate handlers.

There is variation between states in both the number of certificate holders and also the proportion of different types of USDA licenses or registrations. In terms of geographic region, the 12 Midwestern states had 3,408 certificates, followed by 2,457 certificate holders in the 16 Southern states and DC. The Western 13 states and nine states in the Northeast held 1,081 and 1,085 certificates, respectively. Analysis of the factors that affect variation across the states is beyond the scope of this paper. However, it is likely that population, tourism, higher education, and biotechnology play a role. For instance, the overall number of certificate holders is likely associated with state population, the number of exhibitors with tourism, and the number of registered research facilities with the number of biotechnology companies and research institutions. One clear outlier in the state data is Missouri, with a number of facilities – largely breeders – that is many standard deviations higher than other states. The reason for this is not obvious.

Together the data reported here provide an accurate context for better conveying the distribution of settings in which animals covered by the AWA live in the US. The data show that the number of entities follows an orderly pattern with respect to factors associated with the number of type of entities. The picture is necessarily incomplete, however. To provide a more complete picture would require data on animal numbers. Data on AWA-covered species is routinely reported in a consistent and publicly accessible form only for animals in research facilities. It is therefore not possible to determine whether, and how, the number of certificates issued to exhibitors, dealers, and breeders is associated with the number of animals. For instance, it may be the case that the total number of animals in exhibitor-licensed facilities is equal to the number in breeding or research facilities, despite large differences in the total number of certificate holders. Better understanding of the animal numbers and proportion of animals in each setting remains for future analysis but will be important in order to provide a complete picture and context for public consideration and decision-making about practices and policies related to captive animal care.

### Estimates and reports on number of animals

The current report is focused on USDA certificate holders. As highlighted above, however, the broad context for considering the use of animals in captivity in the US includes both animals that fall under the AWA and those that are excluded. Toward that goal, we provide a synopsis here of available information and estimates of the number of animals in various captive settings in the US. In order to provide an equivalent comparison, the focus is largely on those species that fall under the AWA. Fish, amphibians, and reptiles are excluded. Figure 5 provides a basic illustration of the estimated number of animals in three broad categories of use, while Table 2 provides detailed information. It is critical to note the following caveats: While there are federally-reported data or data from trade organizations that estimate the number of animals that are pets, in agriculture, or in research, testing, or education, the number of animals in a range of other activities is not federally tracked or published and thus remains largely unavailable for comparison. For instance, the number of animals involved in exhibition, zoos, and entertainment is not publicly reported. Further, for AWA-species that live in facilities that are not required to have USDA certificate, including sanctuaries, the number of animals is not publicly available. These caveats indicate that the comparisons are estimates. They also point to a gap in public information that results from variation federal in requirements (see Figure 2 above).

**Table 2.**
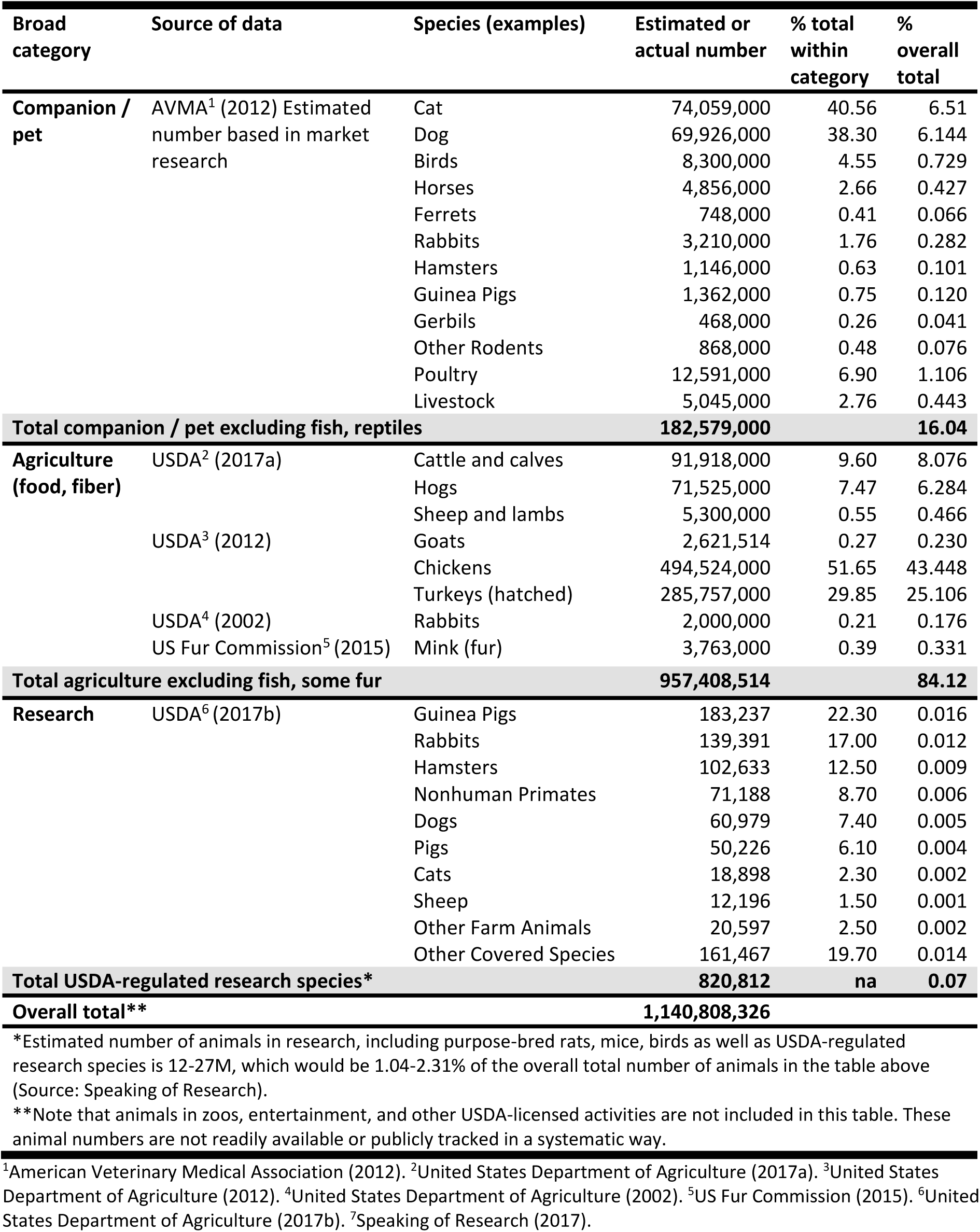
Estimates of number of animals in different activities in US

**Figure 5.**
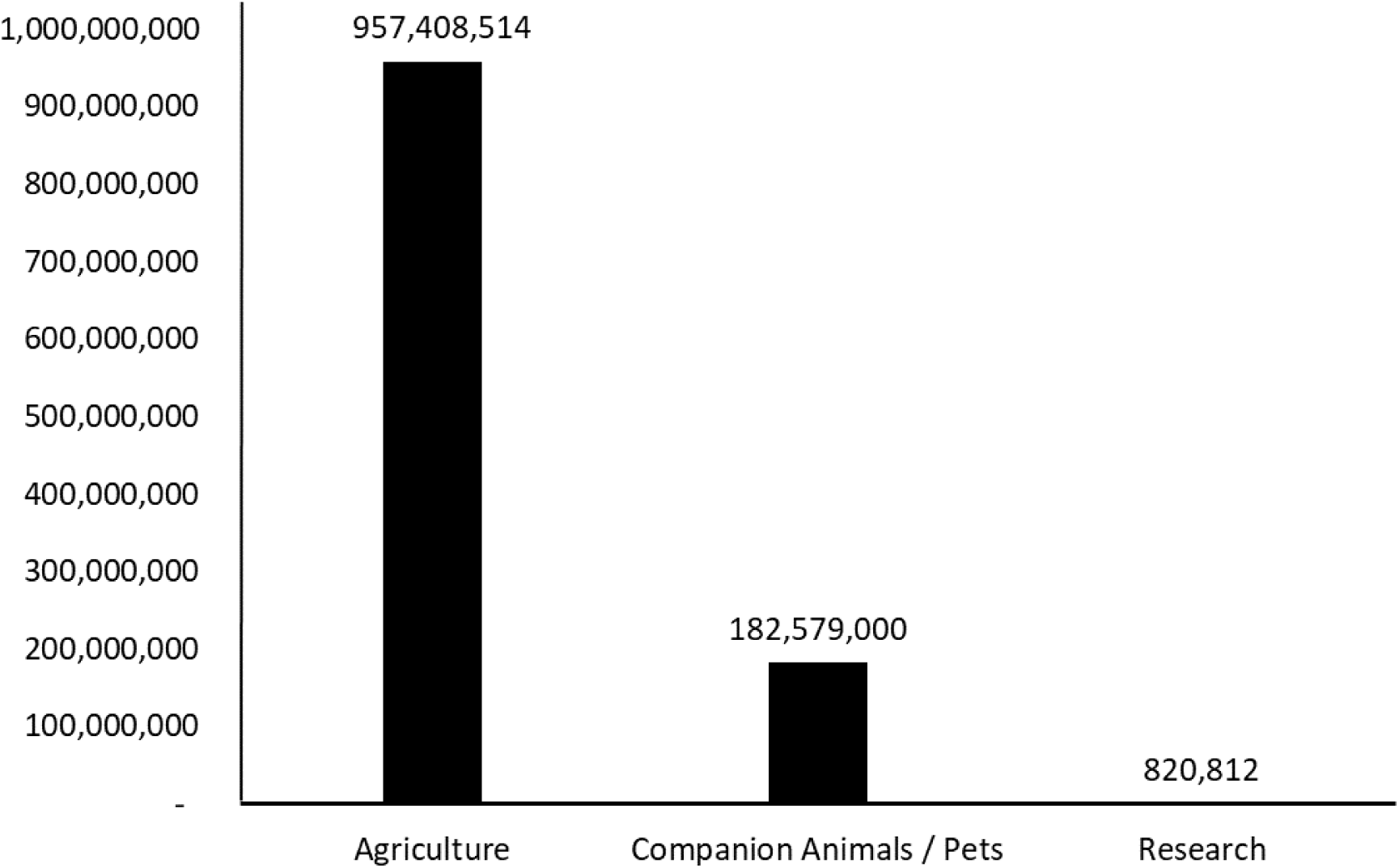
Comparison of the estimated total number of animals in the US in three major activities: agriculture, pets, and research/testing/education.

#### Number of agricultural animals

In brief, in the US, an estimated 957,408,514 animals are used in agriculture as reported by the USDA. Animals counted as agricultural by the USDA are those used as food or fiber. The data in Table 2 include cattle and calves, hogs, sheep and lambs, goats, chickens, turkeys, rabbits, and mink reported in the USDA 2017 Agriculture Statistic Annual Report, the 2012 Census of Agriculture, the US Rabbit Industry Profile, and the US Fur Commission production statistics.^13,16,17,18^ The nearly one billion animals make up roughly 84% of all animals used for agriculture, companionship, and research/testing/education.

#### Number of companion animals

There is no federal requirement for licensure of companion animals, nor centralized public reporting of companion animal numbers in the US. Thus, the real number of animals serving as human companions, or pets, is largely unknown. The American Veterinary Medical Association (AVMA) engages in market research that provides detailed population estimates of companion animal ownership.^1^ The AVMA’s public data includes basic information about the estimated number of a variety of companion animals, including exotic animals. Table 2 shows the number of each type of companion animal (cats, dogs, birds, horses, ferrets, rabbits, hamsters, guinea pigs, gerbils, other rodents, poultry, livestock) reported by the AVMA. According to the AVMA, in the US there are roughly 182,579,000 companion animals of these broad species categories, making up roughly 16% of all animals used for agriculture, companionship, and research/testing/education.

#### Number of animals in research, testing, and education

The number of USDA-covered research animals is one of the most well-documented census numbers for captive animals in the US. As discussed above and highlighted in Figure 2, under the AWA, research facilities are required to report their animal census annually and these reports are made public and easily accessible. Table 2 includes the 2017 USDA annual reports of animal usage by fiscal year to provide the number guinea pigs, rabbits, hamsters, nonhuman primates, dogs, pigs, cats, sheep, other farm animals, and other covered species used in research, testing, or education with research-registered facilities for the year 2016.^14^ The total number of these animals was 820,812, making up 0.07% of all animals used for agriculture, companionship, and research/testing/education.

Rats, mice, and birds bred for research are not subject to the annual USDA report requirement and are estimated to comprise over 95% of animals in research, with an estimated range of 12-27 million used.^7^ For the purpose of comparison and context, Table 2 also shows how using this larger number would affect the estimated percentage of animals in research and testing relative to the proportion in agriculture and the number of pets. Increasing the estimate of animals in research to 12-27 million in order to reflect the number of mice, rats, and birds would raise the overall percentage to between 1.04% and 2.31%. Thus, even with the most liberal estimate, animals in research comprise ∼2%, a number that would decrease further with inclusion of animals in exhibition, entertainment, and other public display.

#### Animals in exhibition

There are few estimates of the overall number of animals in various forms of exhibition and entertainment. Although the great majority of animals housed in these settings fall under the requirement for a USDA license for exhibition, unlike registered research facilities, exhibitors are not mandated to submit an annual report with animal census numbers that are then made publicly available. The number of animals may appear in a facility’s USDA inspection report; however, a review of those reports indicates that the census is often missing. Further, given that routine inspections are not mandated for exhibitors (see Figure 2), there is also no assurance that analysis of publicly-available inspection reports would capture all licensed exhibitors. Overall, there is no direct and straightforward way to assemble a publicly-sourced, reliable, national figure for the number of animals in the wide range of settings that together comprise the USDA exhibitor license category. What is readily available from the USDA website is a complete list of all of the entities that hold USDA licenses and registration. Although these data do not directly address the question of animal numbers, they do provide an accurate view of the broad characteristics of the entities that fall under the AWA.

#### Animals in sanctuary, refuge, and non-display

Under the AWA, a wide variety of entities that house species covered by the law are not required to be licensed. These entities generally include facilities that do not charge the public for entry to view the animals and do not breed the animals for sale. Sanctuaries, some wildlife refuges or rescues, and private zoos may all fall under this category. Although it is not necessarily required, some sanctuaries do apply for USDA certificates as licensed exhibitors. Given the variation and the lack of an animal number reporting requirement, it is not possible to estimate the number of animals in US sanctuaries, refuges, or rescues.

### Concluding remarks

The USDA is charged with oversight of over 8,000 certificate holders, including licensed and registered facilities. The responsibility includes performing the federally-mandated routine annual inspections at 1,106 research facilities. For the greater majority of entities, including 2,628 exhibitors and 2,701 breeders, the USDA is not required to perform annual routine inspections, but must provide oversight that can include inspections every three years or upon receipt of complaints or concerns.^15^ The descriptive data here provide a foundation for better evaluation of the feasibility and potential impact of proposals for changes in USDA oversight of animals housed in a range of captive settings that fall under AWA. For instance, in light of the fact that research facilities comprise less than one-fifth of certificate holders, it is clear that proposals to address the current unevenness in requirements and inspections between research facilities and exhibitors would require substantial changes in USDA resources and personnel. Consideration of the number and proportion of certificate holders provides a more inclusive context for understanding the relative distribution of resources used for oversight. The data and comparisons described here also point to gaps in information that is essential to accurately convey the number of animals used in the broad range of human activities.

## Acknowledgements

We are grateful to Drs. Sangeeta Panicker, Taylor Bennett, and Peter Pierre for helpful discussions of evidence, practices, and policies in captive animal care and oversight that informed preparation of this paper. We also appreciate the efforts of our undergraduate student research team members who provided technical assistance: Trevor Gauthier, Brooke Meidam, and Amanda Novak. This research was partially supported by the University of Wisconsin-Madison Department of Psychology, Graduate School, Wisconsin Alumni Research Foundation.

